# Genetic Analysis and Functional Assessment of a *TGFBR2* Variant in Micrognathia and Cleft Palate

**DOI:** 10.1101/2024.04.08.588524

**Authors:** JES-Rite Michaels, Ammar Husami, Andrew M. Vontell, Samantha A. Brugmann, Rolf W. Stottmann

## Abstract

Cleft lip and cleft palate are among the most common congenital anomalies and are the result of incomplete fusion of embryonic craniofacial processes or palatal shelves, respectively. We know that genetics play a large role in these anomalies but the list of known causal genes is far from complete. As part of a larger sequencing effort of patients with micrognathia and cleft palate we identified a candidate variant in *transforming growth factor beta receptor 2* (*TGFBR2*) which is rare, changing a highly conserved amino acid, and predicted to be pathogenic by a number of metrics. The family history and population genetics would suggest this specific variant would be incompletely penetrant, but this gene has been convincingly implicated in craniofacial development. In order to test the hypothesis this might be a causal variant, we used genome editing to create the orthologous variant in a new mouse model. Surprisingly, *Tgfbr2*^*V387M*^ mice did not exhibit craniofacial anomalies or have reduced survival suggesting this is, in fact, not a causal variant for cleft palate/ micrognathia. The discrepancy between in silico predictions and mouse phenotypes highlights the complexity of translating human genetic findings to mouse models. We expect these findings will aid in interpretation of future variants seen in *TGFBR2* from ongoing sequencing of patients with congenital craniofacial anomalies.

## Introduction

Cleft lip/cleft palate are craniofacial congenital anomalies characterized by incomplete fusion of the lip and/or palate during embryonic development. These are the among the most common of all congenital anomalies with an incidence of approximately 1/700 live births and (Mossey et al., 2009; Murray, 2002). Causes of cleft lip and cleft palate are quite diverse with known contributions from genetics and environmental influences (teratogens) as well as the interplay between each (Beames and Lipinski, 2020; Dixon et al., 2011). Genetic studies of orofacial clefting disorders in humans have identified a handful of genes known to have high association with variable forms of cleft lip/ palate such as *IRF6* (Schutte et al., 1993), *FOXE1* (Moreno et al., 2009), and *TP63* (Sutton and van Bokhoven, 1993). These studies to date have been plagued with incomplete penetrance complicating a complete understanding of cleft lip/palate genetics (Leslie, 2022; Reynolds et al., 2020). We have been sequencing families with syndromic cleft lip/palate but no known genetic diagnosis to continue to build our knowledge of genetic variants leading to these conditions and potentially design future treatments.

Transforming Growth Factor (TGF) signaling is a classical developmental signaling pathway with extracellular ligands binding to complexes of transmembrane receptor tyrosine kinases. Ligand binding facilitates phosphorylation of Type II receptors which then activates intracellular Smad proteins to ultimately traffic to the nucleus and facilitate transcription of direct targets (Massague, 2012; Ross and Hill, 2008; Schmierer and Hill, 2007; Shi and Massague, 2003). TGF-β signaling mediates several key functions during embryonic development including cell proliferation, differentiation, and extracellular matrix formation (Chai et al., 2003). TGF superfamily signaling has strongly been associated with craniofacial development(Iwata et al., 2011; Ueharu and Mishina, 2023). While the *Tgfbr2* null embryos die by E10.5 (Oshima et al., 1996), the role of *Tgfbr2* in craniofacial development became clear with studies using conditional inactivation in the cranial neural crest cells using the *Wnt1Cre* driver. This ablation of *Tgfbr2* result in cleft palate due to reduced cell proliferation in the palatal mesenchyme at E14.5 (Ito et al., 2003; Iwata et al., 2012a; Iwata et al., 2011; Iwata et al., 2014a; Iwata et al., 2014b; Iwata et al., 2012b). In addition, these neural crest specific *Tgfbr2* mutants exhibited smaller mandible with reduced condylar, coronoid process and a complete lack of the angular process of the mandible in the proximal region (Ito et al., 2003; Oka et al., 2007). Here we identify a candidate variant in *transforming growth factor beta receptor 2* (*TGFBR2*). To directly test this hypothesis this may be a new causal variant for isolated micrognathia and cleft palate, we utilize the power of mouse genetics and precision genome editing.

## Results

### A potentially pathogenic TGFBR2 variant involved in craniofacial anomalies

We performed whole genome sequencing as part of a larger project on the human genetics of syndromic cleft lip/palate phenotypes. This analysis identified a candidate variant in *transforming growth factor beta receptor 2* (*TGFBR2*, NCBI Gene ID: 7048). This variant (NM_003242.5:c.1159G>A; Chr3(GRCh37):g.30713824G>A; p.Val387Met) changes the amino acid at position 387 from a valine to a methionine. This variant has been noted previously (rs35766612) and is present at extremely low levels in control population databases with an overall allele frequency of 0.0017 and 4 homozygotes in gnomAD v4 (Chen et al., 2024), a virtually identical allele frequency of 0.0017 in the Regeneron million exome collection (Sun et al., 2023), and a combined annotation dependent depletion (CADD: (Rentzsch et al., 2019) score of 24.4. The nucleotide sequence in this region of the genome is moderately conserved across phylogeny, but the amino acid code is faithfully conserved through *C. elegans*, with the exception of the *D. melanogaster* genome (Fig. 1A). The change from Valine to Methionine only has a small physicochemical distance with a Grantham score of 21 (0-215) but is in the protein kinase domain which is critical for TGFBR2 function (Lin et al., 1992). The variant is predicted to be “deleterious” by SIFT (Ng and Henikoff, 2003) (score = 0), “disease causing” by MutationTaster (Schwarz et al., 2014) (prob = 1), “probably damaging” by PolyPhen-2 (Adzhubei et al., 2013) (score = 0.996), and “intolerant” by MetaDome (Wiel et al., 2019) (score = 0.27). Previous reports in ClinVar (Landrum et al., 2014) have conflicting interpretations of pathogenicity but were evaluated in different pathological contexts such as Marfan syndrome, Loeys-Dietz syndrome and aortic disease. We therefore hypothesized that this variant may be a cause of penetrant cleft palate and micrognathia in humans.

**Figure 1.**
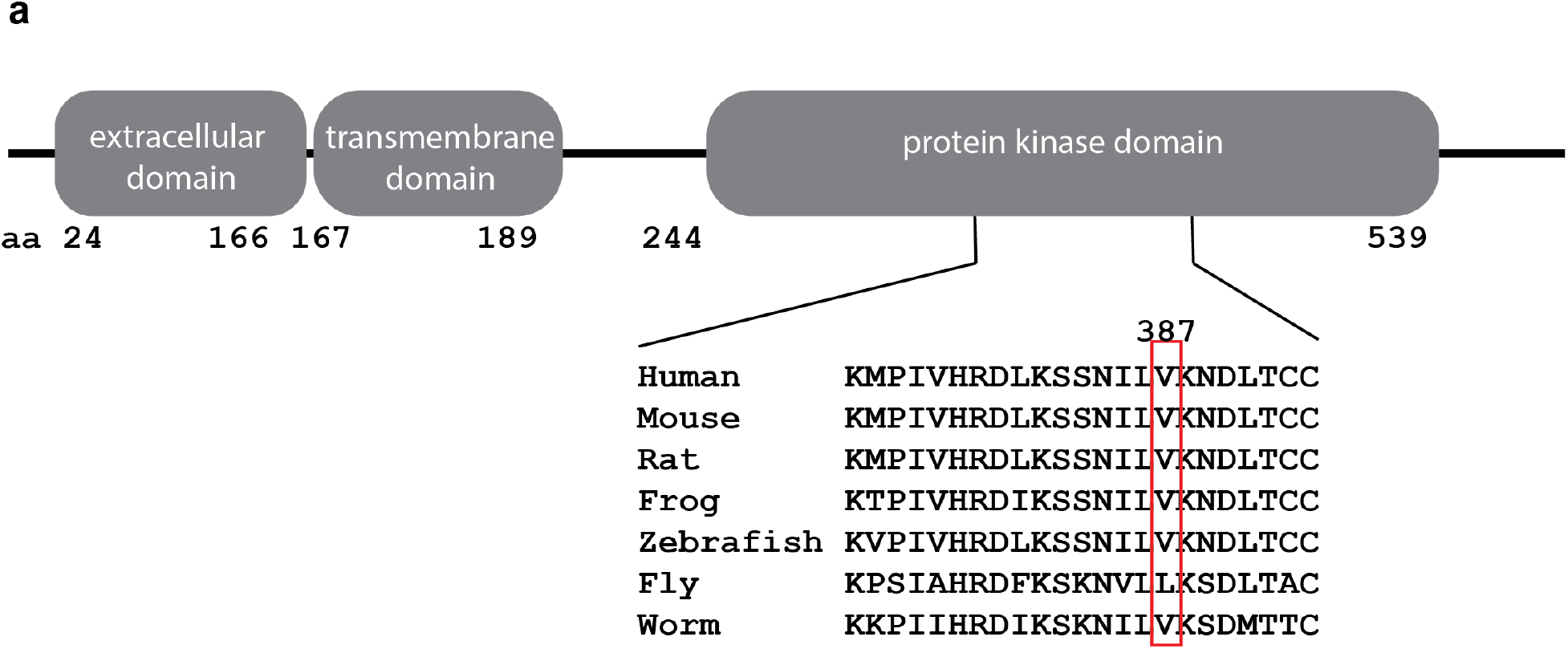
A novel TGFBR2 allele. (A) TGFBR2 protein domain structure. V387 is in an area of highly conserved sequence in the kinase domain.

### A novel mouse model of Tgfbr2^V387M^

Given the known role for *Tgfbr2* in craniofacial development and the incompletely penetrant nature of many craniofacial variants in human, we created a mouse model to test the pathogenicity of this missense variant. This portion of the TGFBR2 protein is highly conserved between human (Fig. 2A) and mouse (Fig. 2B). CRISPR-CAS9 genome editing was used in one cell mouse embryos with injection of a mixture containing *Cas9*-gRNA and both variant knock-in and silent wild-type donors. Multiple founders were identified with the desired knock-in with Sanger sequencing and were further bred with wild-type females. Animals from this outcross and their progeny were used in the analysis reported here. We designate this allele as *Tgfbr2*^*em1Rstot*^ but refer to it hereafter as *Tgfbr2*^*V387M*^. Genotyping was performed with a combination of Sanger sequencing (mutant allele shown in Fig. 2C) and PCR followed by restriction digest as a variant in the donor oligonucleotide created a BspH1 recognition site (Fig. 2D).

**Figure 2.**
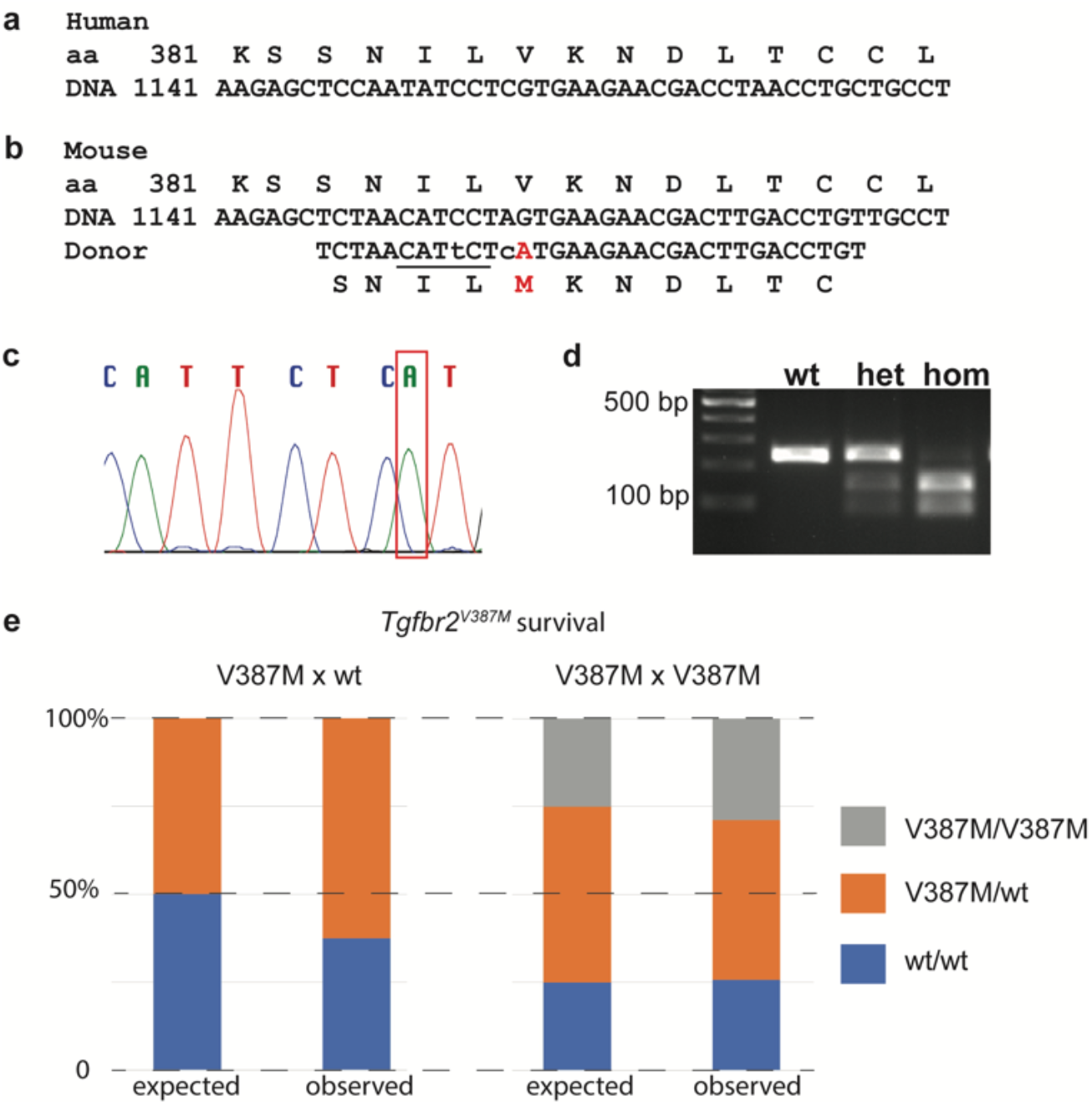
A mouse model of Tgfbr2^V387M^ missense allele. Amino acid and DNA sequence of *Tgfbr2* shows high conservation between human (A) and mouse (B). (C) Sanger sequence of mice showing desired sequence change from G to A indicated by the red box in mutants. (D) PCR genotyping followed by restriction digest indicating ability to clearly differentiate wild-type, heterozygous, and homozygous mutant mice. (E) Survival of all mice are not significantly different than Mendelian expectations.

### Tgfbr2^V387M^ variant does not cause any craniofacial phenotypes

We first analyzed survival of *Tgfbr2*^*V387M/wt*^ heterozygotes from crosses with wild-type animals and see no reduced recovery of heterozygotes (n=20 from 32 total animals; Chi-squared p-value = 0.16; Fig. 2E, Table 1). To test if homozygotes have a phenotype, we intercrossed heterozygotes and saw no deviation from expected ratios of *Tgfbr2*^*V387M/wt*^ heterozygotes or *Tgfbr2*^*V387M/V387M*^ homozygotes from 66 weaned animals (Chi-squared p-value = 0.72; Fig. 2E, Table 1). We never noted cleft palate during observation of any animals during the course of this study. Given that cleft palate is perinatal lethal in mouse models, we interpret these findings to suggest that cleft palate was very rarely, if ever, present in animals carrying this predicted pathogenic variant at a highly conserved position. One mechanism potentially leading to cleft palate is micrognathia in which the hypoplastic jaw leads to a crowding of the oral cavity and the posteriorly displaced tongue interferes with palatal shelf elevation and/or closure (Diewert, 1986). While this was not severe enough to cause cleft palate in these animals if present at all, we tested if mutants may have micrognathia as suggested by the human participants. We performed skeletal preparations of animals born from *Tgfbr2*^*V387M/wt*^ heterozygous intercrosses to highlight cartilage and bone and measured the length of the mandible as well as the total head length (Fig. 3A,B). Analysis did not show any reduction between genotypes in mandible length as an isolated measurement or relative to skull length, either at postnatal day (P) 60 (Fig. 3G,H, n=50) or P120 (Fig. 3G,H, n=24). We also examined the roof of the mouth for any subtle deficits and noted no abnormalities (Fig. 3C,D). Finally, we performed histological analysis of a number of *Tgfbr2*^*V387M/V387M*^ homozygous mutants at E18.5 and P0 and saw no subtle craniofacial phenotypes in these animals either (Fig. 3E,F). We conclude from these data that the *Tgfbr2*^*V387M*^ missense variant does not lead to any craniofacial phenotypes in the mouse.

**Table 1.** Survival of *Tgfbr2*^*V387M*^ animals at weaning.

|  | Total | wt/wt | V387M/ wt | V387M/V387M | Chi-sq. |
| --- | --- | --- | --- | --- | --- |
| **V387M x wt** |  |  |  |  |  |
| expected | 32 | 16 | 16 | - |  |
| observed | 32 | 12 | 20 | - | 0.157 |
| **V387M x V387M** |  |  |  |  |  |
| expected | 66 | 16.5 | 33 | 16.5 |  |
| observed | 66 | 17 | 30 | 19 | 0.716 |

**Figure 3.**
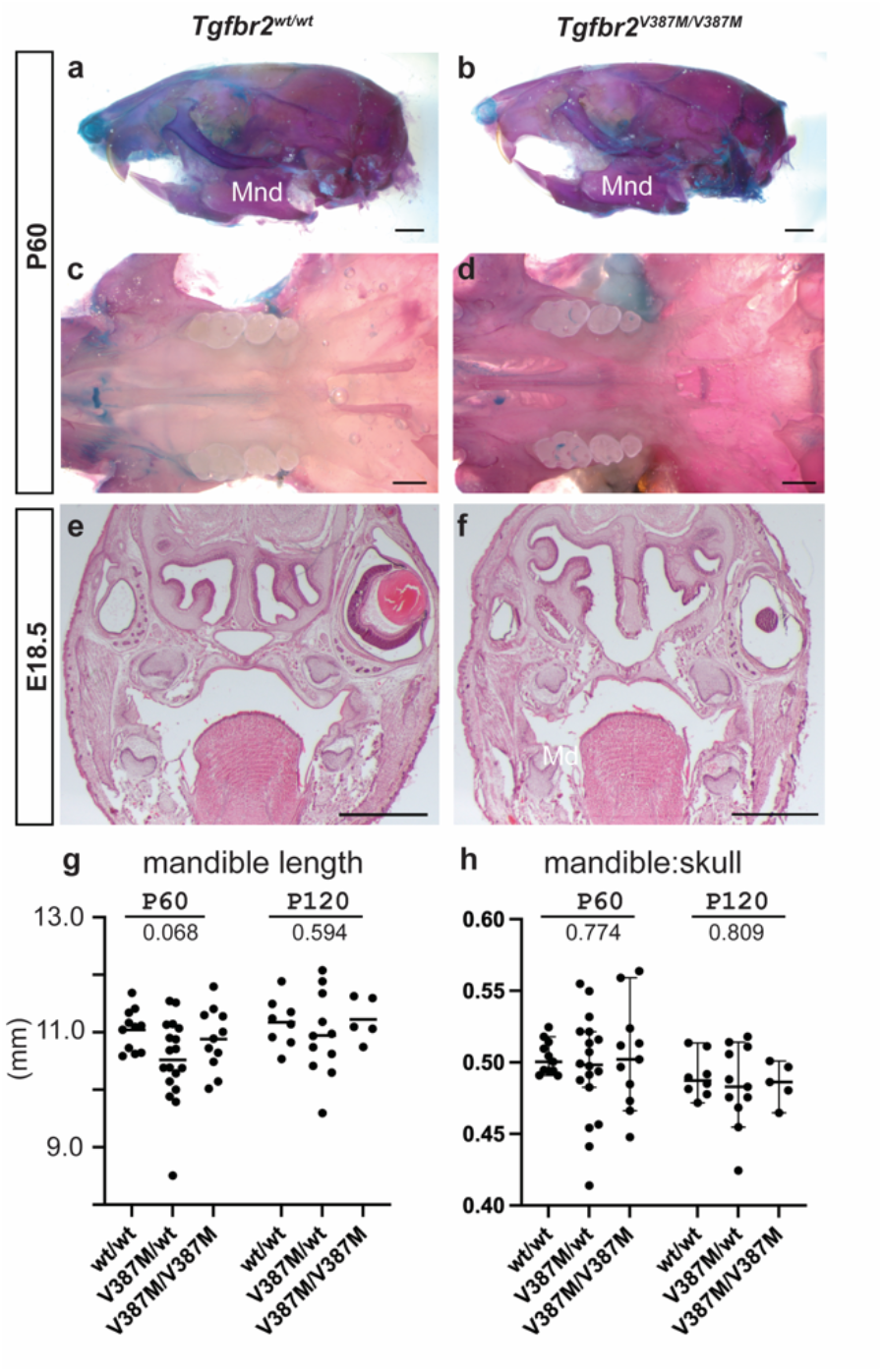
Craniofacial analysis of Tgfbr2^V387M^ mice. (A-D) Skeletal preparations of animals at postnatal day (P) 60 from the lateral view (A,B) or focused on the palatal surface (C,D) do not appear any different between wild-type (A,C) and *Tgfbr2* ^*V387M/V387M*^ homozygous mutants (B,D). (E,F) Histological analysis of wild-types (E) and mutants (F) at E18.5 also did not reveal any differences. (G,H) Mandible length was measured as an isolated element (G) or as a ratio to total skull length and shows no difference between genotypes at P60 or P120. Scale bars in A-F indicate 1mm. Statistical values shown are from an ANOVA.

## Discussion

Here we present a novel missense variant mouse allele of *Tgfbr2* created to assess the hypothesis the variant is causing disrupted craniofacial development with cleft palate due to micrognathia (Pierre Robin Sequence). While this variant was predicted to be deleterious by multiple algorithms, our findings in the mouse do not support a biological role for this variant as we see no reduced survival of *Tgfbr2*^*V387M/wt*^ heterozygous or *Tgfbr2*^*V387M/V387M*^ homozygous animals and do not find any phenotypes reminiscent of human participants. We conclude this is not likely to be a causal variant in human genetics, but recognize this might be a failure of the mouse model to accurately recapitulate human biology.

*TGFBR2 has known roles in human biology* Heterozygous missense variants in *TGFBR2* are well established to cause Loeys-Dietz Syndrome (Loeys et al., 2005; Loeys and Dietz, 1993). While cleft palate can be seen in some of these patients (two of the first six *TGFBR2* participants reported), the more common clinical characteristics include arterial aneurysm/dissection, skeletal phenotypes such as scoliosis or joint laxity, and craniosynostosis. The patients reported here were not known to have any of the clinical characteristics of Loeys-Dietz Syndrome and this particular variant we find has been most often classified as (likely) benign when analyzed in this context by the clinical community.

### Tgfbr2 in craniofacial biology

*Tgfbr2* has been deleted in the mouse and homozygous null embryos die by E10.5 (Oshima et al., 1996). A *Wnt1*-Cre mediated deletion in neural crest cells was much more informative (Ito et al., 2003). These mutants have a completely penetrant complete failure of fusion in the secondary palate, likely due to reduced proliferation of the neural crest cells as they facilitate horizontal growth of the palatal shelves during mid-gestation (E14.5)(Ito et al., 2003; Iwata et al., 2012a; Iwata et al., 2011; Iwata et al., 2014a; Iwata et al., 2014b; Iwata et al., 2012b). In addition, CNCC-specific inactivation of *Tgfbr2* resulted in a hypoplastic mandible with severe defects in the proximal region of the mandible (Ito et al., 2003; Oka et al., 2007). This conditional ablation of *Tgfbr2* also leads to skull defects which are also seen in Loeys-Dietz Syndrome patients. Neither the mouse model we report here, nor the sequenced family show any evidence for craniosynostosis or defects of the calvarial bones.

### Mouse as a model for human congenital craniofacial anomalies

The genetic similarities, experimental tools and overall conserved processes of development and disease between mouse and humans have supported the rise of the mouse as the most powerful mammalian model of human development and disease. The power in this approach has only grown with the advent of CRISPR-CAS9 mediated genome editing and the ability to manipulate the genome to model specific variants seen in human. This is the approach we took here with the aim of assessing pathogenicity of a specific variant identified through whole genome sequencing. In addition to addressing specific hypotheses about genetic causes of human disease, these mouse models can facilitate experiments to determine underlying molecular mechanism(s) and potentially test proof of concept experiments for therapeutic interventions.

One alternative explanation for our data is that the *Tgfbr2* variant a risk factor in humans, but this is not recapitulated in the mouse due to subtle differences in the two models. While the processes of craniofacial development broadly, and palatogenesis specifically, are quite similar, there are some developmental differences (Gritli-Linde, 2008; Yu et al., 2017). In general, however, there is good concordance between the mouse and human model to date. A recent review compiled a list of mouse models with cleft palate and human disorders associated with those genes (Funato et al., 2015). In this comparison, 15 human diseases with cleft palate as a phenotypic feature had a corresponding mouse models cataloged and all were noted to have cleft palate, including the *Tgfb2* ligand itself (Sanford et al., 1997). Another review found a similarly very high concordance (Gritli-Linde, 2008). Even in the biochemical pathway we are considering here, we have mentioned multiple mutants of the TGFB signaling cascade which have cleft palate in mouse mutants (e.g., (Ito et al., 2003; Sanford et al., 1997). Another variable to consider is the genetic background. We conducted our experiments only with mice on a C57BL/6J background. Mouse strains are known to have different sensitivities, including environmental influences, to clefting (Biddle and Fraser, 1976) but the C57BL/6J mouse has not been previously shown to be particularly refractory to craniofacial differences. Indeed, a classic mouse model of Treacher Collins Syndrome is more acutely affected in C57BL/6J as compared to the DBA strain (Dixon et al., 2006). We therefore acknowledge this as a caveat to the data but find it more likely that this *Tgfbr2* variant is not a causal allele for cleft palate or micrognathia leading to palate disruption in mammals despite the deep conservation and *in silico* predicted pathogenicity. In this case, the population data was not consistent with a highly penetrant phenotype. These population frequency datasets are getting more powerful with the addition of new data and will therefore become increasingly valuable and predictive in future analyses. We recognize that periodic reanalysis of exome and genome data has been shown to be productive in solving cases (Hills et al., 2023) and look forward to further hypotheses for the participants described here.

## Methods

### DNA Sequencing

DNA was collected as part of an IRB-approved recruitment protocol and purified using the Qiagen DNeasy Kit and manufacturer’s protocols (Qiagen, USA). Whole genome sequencing was done at Novogene USA to an average depth of at least 30x with standard protocols.

### Variant Discovery

VCF format small SNP and InDels were annotated and filtered using standardized protocols using GoldenHelix VarSeq and Qiagen - Qiagen Clinical Interpreter – Translational (formerly Ingenuity) variant analysis software packages. The variant annotation and interpretation analyses were generated through the use of Ingenuity® Variant Analysis™ software https://wwusingtics.com/products/ingenuity-variant-analysis from QIAGEN, Inc. Variants were filtered based on population frequencies. Subsequently, variants known or predicted to have semantic similarity with Micrognathia, and cleft palate were discovered using Phenotype Driven Ranking (PDR) filter.

### Mouse model creation

Mice (NCBI Taxon ID 10090) were created at the Cincinnati Children’s transgenic animal and genome editing core (RRID:SCR_022642). Mouse zygotes (C57BL/6N strain) were injected with 200 ng/μl CAS9 protein (IDT and ThermoFisher), 100 ng/μl *Tgfbr2*-specific sgRNA (AGGTCAAGTCGTTCTTCACT), 75 ng/μl single-stranded donor oligo-nucleotide (KI) to create the *Tgfbr2* variant (AGAGCTGGGCAAGCAGTACTGGCTGATCACGGCGTTCCACGCGAAGGGCAACCTGCAGGAGTACCTCACGAGGCATGTCATCAGCTGGGAGG ACCTGAGGAAGCTGGGCAGCTCCCTGGCCCGGGGATCGCTCATCTCCACAGTGACCACACTCCTTGTGGGAGGCC; IDT, Iowa) and 75 ng/μl single-stranded donor oligo-nucleotide of wild-type sequence (WT-S) with silent variants (TGTTGGCCAGGTCATCCACAGACAGAGTAGGGTCCAGGCGCAAGGACAGCCCGAAGTCACACAGGCAACAGGTCAAGTCGTTCTTCAcgAGa ATGTTAGAGCTCTTGAGGTCCCTGTGAACAATGGGCATCTTG; IDT, Iowa)followed by surgical implantation into pseudo-pregnant female (CD-1 strain) mice. Both donors are used together to prevent possible lethality of homozygous target mutations. Silent mutations (indicated by the lowercase letters in Fig. 2B) are also introduced in the knock-in donor to prevent recutting of the target allele by CRISPR and to create a restriction enzyme site to facilitate genotyping. PCR genotyping was performed by amplification of genomic DNA (F:CATCGCTCATCTCCACAGTGAC and R:TGAAGCCAGGCATGAAGTCTGAG primers).

The PCR products were subject to digestion with BspH1 or Sanger Sequencing and pups exhibiting editing of interest were then crossed to wild-type (C57BL/6J) mice and the resulting progeny were Sanger sequenced (CCHMC DNA Sequencing and Genotyping Core) to confirm the alleles generated. Propagation of the *Tgfbr2* allele was done by crossing to wild-type C57BL/6J mice and/or intercross. Further genotyping was with a combination of PCR followed by restriction digest and/or Sanger sequencing.

### Animal housing

All experiments using mice in this study were performed using ethically acceptable procedures as approved by the Institutional Animal Care and Use Committee at Nationwide Children’s Hospital (AR21-00067). Mice were fed mouse breeder diet and housed in ventilated cages with a 12 h light/12 h dark cycle.

### Skeletal preps and pictures

Skeletons from P60 or P120 animals were stained for Alcian Blue and Alizarin Red to visualize cartilage and bone, respectively. Briefly, animals were skinned and eviscerated and fixed for 2 days in 95% ethanol. They were stained overnight at room temperature in 0.03% (w/v) Alcian Blue solution (Sigma-Aldrich, A3157) containing 80% ethanol and 20% glacial acetic acid. Samples were destained in 95% ethanol for 24 h followed by pre-clearing in 1% KOH overnight at room temperature. Skeletons were then stained overnight in 0.005% Alizarin Red solution (Sigma-Aldrich, A5533) containing 1% KOH. A second round of clearing was performed by incubating tissues in 20% glycerol/1% KOH solution for 24 h. Finally, they were transferred to 50% glycerol/50% ethanol for photography. Skeletal preparations were imaged using a Zeiss Discovery.V12 Stereoscope and length measurements were recorded for mandibular bones and skull length (tip of nasal bone to basoccipital bone).

### Histology

Hematoxylin & Eosin (H&E) fixation was in Bouin’s solution followed by washes in 70% ethanol. For H&E staining, embryos were embedded in paraffin and cut to 10 μm sections before staining using standard techniques (Behringer, 2014). All images were taken via Zeiss Discovery.V12 Stereoscope. Paired images are shown at the same magnification.

### Statistics

All statistical analyses were performed using GraphPad Prism 9.5.1. Ordinary one-way ANOVA with Dunnett’s post-hoc multiple comparisons test was performed for comparison of mandible lengths. ANOVA P values are indicated on all graphs. The data shown are the mean ± 95% confidence interval. No samples were excluded from analyses.

## Acknowledgements

We appreciate comments on the manuscript from the Stottmann laboratory group. Funding for this project comes from R35DE027557 (S.A.B.) and R01DE027091 (R.W.S.). The Regeneron Genetics Center, and its collaborators (collectively, the “Collaborators”) bear no responsibility for the analyses or interpretations of the data presented here. Any opinions, insights, or conclusions presented herein are those of the authors and not of the Collaborators. This research has been conducted using the UK Biobank Resource under application number 26041.

